# Neonatal LPS exposure reduces ATP8A2 level in the prefrontal cortex in mice via increasing IFN-γ level

**DOI:** 10.1101/2020.08.09.243477

**Authors:** Jiapeng Deng, Linyang Song, Zhiqin Yang, Sixie Zheng, Zhuolin Du, Li Luo, Jing Liu, Xiaobao Jin, Junhua Yang

## Abstract

Neonatal lipopolysaccharide (LPS) exposure can cause depressive-like behaviors in rodents involving elevated interferon (IFN)-γ. Studies have linked down-regulation of prefrontal cortex (PFC) ATPase phospholipid transporting 8A2(ATP8A2) expression to depressive-like behaviors. In non-neuronal cells, IFN-γ could reduce ATP8A2 expression. We therefore hypothesized that neonatal LPS exposure might induce PFC ATP8A2 down-regulation by increasing IFN-γ level. Here, C57BL6/J mice of both sexes received 3-dose-injections of LPS (50μg/kg bodyweight, i.p.) on postnatal day (PND)5, PND7 and PND9. LPS-treated mice showed a transiently decreased PFC ATP8A2 expression. Moreover, a negative correlation of PFC ATP8A2 expression was found with IFN-γ level. Using neutralizing mAb, IFN-γ was identified as the key mediator of LPS-induced PFC ATP8A2 decrease. Besides, neutralizing IFN-γ during neonatal LPS exposure attenuated the depressive-like behaviors in adulthood. In sum, neonatal LPS exposure reduced ATP8A2 level in PFC in mice via increasing IFN-γ level, maybe associated with mechanism underlying LPS-induced brain and behavior impairments.

## Introduction

ATP8A2, a member of the P4-ATPase family in mammals, is highly expressed in the brain, spinal cord, testis, and retina, which flipping phosphatidylserine and phosphatidylethanolamine across the cell membrane (Andersen et al.,2016). It plays an important role in maintaining the stability and normal function of the cell membrane (Coleman et al.,2009). Therefore, ATP8A2 in the brain is vital for normal neural development and function (Coleman and Molday,2011; Xu et al.,2012), since these processes involve cell proliferation, migration, synapses pruning of neurons (Brandon and Sawa,2011). ATP8A2 expressed in the brain has received extensive attention recently and researches have shown that reduced expression of ATP8A2 leads to axonal mutation and neurodegenerative diseases (Choi et al.,2019; Zhu et al.,2012). There is also research reporting that ATP8A2 expression decreases in conditions of Alzheimer’s disease and bacterial infection (Aaron et al.,2018; Ross et al.,2011). Moreover, depressive-like behavior has been reported to be associated with decreased ATP8A2 expression in the prefrontal cortex (PFC) (Chen et al.,2017). Although ATP8A2 has been widely studied, little is known about how ATP8A2 expression is regulated.

Recently, IFN-γ was shown to significantly reduce the expression of ATP8A2 in non-neuron cells (Shulzhenko et al.,2018). IFN-γ produced in the periphery can penetrate brain parenchyma across the immature blood-brain barrier and even increase the permeability of the mature blood-brain barrier (Hansen-Pupp et al.,2005; Wong et al.,2004). IFN-γ may mediate the neonatal lipopolysaccharide (LPS) exposure-induced depressive-like behaviors in mice (Campos et al.,2014), which is associated with the function of PFC. Besides, IFN-γ expression in PFC increased more than twice after LPS treatment (Laumet et al.,2018). Based on the background mentioned above, it was postulated that neonatal LPS exposure might induce ATP8A2 down-regulation in PFC in mice by increasing the IFN-γ level. There is yet no report addressing this issue.

To test this hypothesis, we challenged neonatal mice with LPS and observed a transiently decreased expression of ATP8A2 in PFC. Using a series of experiments, we further identified IFN-γ as the key mediator of LPS-induced ATP8A2 down-regulation in PFC in mice. These findings may reveal a potential mechanism by which early inflammation leads to impairment of the central nervous system development and function.

## 2. Materials and methods

### 2.1. Animals designing

Litters of newborn C57BL6/J mice (one-day-old) consisting approximately evenly of both sexes were ordered together with their mothers from Guangdong Medical Laboratory Animal Center (Guangzhou, China). They were housed in a specific pathogen-free condition under 12 h light-12 h dark conditions with food and water available ad libitum.

In this study, a randomized block design was used to minimize the effects of systematic error according to publications by statisticians who stated that if the experimenters aim to focus exclusively on the differences between/among different treatment conditions, the effects due to variations between the different blocks should be eliminated by using a randomized block design ANOVA (Refer to https://doi.org/10.1007/978-0-387-32833-1_344;http://www.r-tutor.com/elementary-statistics/analysis-variance/randomized-block-design). In such design, there is only one primary factor under consideration in the experiment. Similar test subjects are grouped into blocks. Then, subjects within each block are randomly assigned to treatment conditions. Each block is tested against all treatment levels of the primary factor at random order. Thus, possible influence by other extraneous factors will by eliminated.

To be specific, a randomized block design was conducted for each of the experiments for investigating potential differences between or among treatment conditions. Each of the blocks consisted of two (for experiments reported in Fig.1 and Fig.2), three (for experiment reported in Fig.5) or five (for experiments reported in Fig.4 and Fig.6) same-sex pups from a same dam. The pups in each block were randomly assigned to different treatment conditions (only one pup in each block for receiving each kind of the different treatment conditions). The sex of n pups in different blocks were same or different, while the litter background of pups in different blocks were certainly different The detailed information indicating how many litters were used and how many pups from each litter were included in each group of each experiment in this study were shown in Supplementary talbe1 to table6 (Supplementary Material). Pups subjected to behavioral tests were weaned on postnatal day (PND)21. Pups subjected to other tests were sacrificed on PND11. This study was approved by the Institutional Animal Ethics Committee of Guangdong Pharmaceutical University.

### 2.2. LPS treatment

Mice were intraperitoneally injected with 3 doses of LPS dissolved in 0.1 mol phosphate balanced solution (PBS) (Escherichia coli O111: B4; Sigma-Aldrich) (50μg/kg bodyweight for each dose;) on PND5, PND7 and PND9. The dosage and schedule for LPS injection were determined based on previous studies (Dinel et al.,2014; Doosti et al.,2013; Liang et al.,2019). The experiment was controlled by intraperitoneal injection of PBS. Littermates were toe-clipped for identification randomly assigned to each of the groups in each experiment as described in Results.

### 2.3 Administration of anti-IFN-γ neutralizing mAb and an isotype IgG1

For experiments conducted in this study as reported in Section 3.3 and Fig.4 to Fig.6, anti-IFN-γ neutralizing mAb and/or an isotype IgG1 (Invitrogen, Thermo Fisher Scientific, Waltham, MA, USA) were intraperitoneally injected daily to mice from PND5 to PND10. By a special experiment, the optimal dosage was first determined as 0.6 mg/kg body weight of anti-IFN-γ neutralizing mAb. The dosage of isotype IgG1 was determined as 0.6 mg/kg body weight accordingly.

### 2.4 Western blot

After being over-anesthetized with 10% chloral hydrate, the mice were decapitated and killed. The mouse PFC tissues were immediately taken on ice, and then homogenized in RIPA lysate containing protease inhibitors (Beyotime, Wuhan, China). After centrifuged at 13000g for 30 min at 4°C, the supernatant was taken. The BCA protein analysis method was used to determine the total protein content of the sample. After mixed with 5×SDS-PAGE loading buffer at the volume ratio of V (protein sample solution): V (5×SDS-PAGE Loading Buffer) = 4:1, the samples were boiled and denatured for 5 min, save for later use. The samples of each group were subjected to SDS polyacrylamide gel electrophoresis. The gel used here is a simplified hand-poured gradient gel, in which the ratio was 40% of the volume being 6% acrylamide and 60% of the volume being 12% acrylamide (using an SDS-PAGE Gel Quick Preparation Kit, Beyotime, Shanghai, China). The procedure to make this gradient gel was according to a previous study (Miller et al.,2016) with modification with respect to the concentrations of acrylamide. When proteins were transferred to PVDF membranes, the membranes were cut according to the indication of the color marker (10-180 kDa, ThermoFisher, Shanghai, China). Two parts of each membrane, one located in the marker’s range 70-180 kDa and the other in the range 25-55 kDa, containing the target proteins ATP8A2 (129 kDa) and the internal control proteins β-actin (43 kDa) were subjected to the subsequent treatment. These membranes were blocked with 5% skimmed milk powder at RT for 2 h. After this, these membranes were put into the corresponding primary antibody incubation solutions, one containing anti-ATP8A2 antibodies (1:500, Abcam, Cambridge, MA, USA) and the other containing anti-β-actin antibodies (1:5000, Abcam, Cambridge, MA, USA) and incubated at 4°C overnight. After rinsing with TBST, these membranes were put into the corresponding secondary antibodies incubation solutions, one containing horseradish peroxidase (HRP)-conjugated goat anti-rabbit antibodies (1:5000, Bioworld, Atlanta, GA, USA) and the other containing HRP-conjugated goat anti-rabbit antibodies (1:5000, Bioworld, Atlanta, GA, USA) and incubated at RT for 1 h. After rinsing with TBST, ECL luminescent solution is added on the membranes in a dark room for exposure and development. Chemiluminescent images were obtained with Carestream XBT X-ray Film (Rayco, Xiamen, Fujian, China), and subsequently scanned and quantified by densitometry using ImageJ software.

### 2.5 Determination of cytokines levels

Forty-eight h after the last LPS injection, the mice were anesthetized deeply with 10% chloral hydrate before the blood was collected from the heart immediately. After blood collection, it was allowed to stand at room temperature for 1 h, and after centrifugation, the supernatant was taken and stored at −70°C for the experiment. Then the mice were transcranial perfused with 0.9% NaCl and PFC was collected immediately on ice. A mouse cytokine/chemokine magnetic bead panel (MCYTOMAG-70K-06; Millipore, Billerica, MA, USA) was employed to detect the levels of IFN-γ, tumor necrosis factor (TNF)-α, interleukin(IL)-1β, IL-6 both in the serum and PFC (Yang et al.,2016). Each of the serum samples was diluted at 1:2 in assay buffer. PFC tissue was prepared as homogenates before assayed. The total protein concentration of each sample was adjusted to 4.5 mg/ml using a BCA protein assay kit (Beyotime, Shanghai, China). Then, the prepared serum and PFC samples were used strictly according to the manufacturer’s protocols for the multiplex assays. The data were collected on a Bio-Plex-200 system (Bio-Rad, Hercules, CA, USA) and analyzed using professional software (Bio-Plex Manager).

### 2.6 Immunofluorescence and cell quantification

Forty-eight h after the last LPS injection, the mice were over-anesthetized with 10% chloral hydrate and transcranial perfused with 0.9% NaCl followed by 4% paraformaldehyde (PFA). After removed, the brains were immediately post-fixed in 4% PFA overnight at 4 °C. Then, the brains were gradient dehydrated with 10%, 20%, and 30% sucrose for 24 h each at 4°C. Free-floating, serial coronal sections (40 μm) were collected on a Leica SM2000R freezing microtome (Leica Microsystems, Richmond Hill, Ontario, Canada) and stored at 4°C for immunostaining. Sections were washed in PBS three times and then blocked in PBS containing 1% bovine serum albumin (BSA) and 0.25% Triton X-100 (Sigma-Aldrich, St. Louis, MO, USA) for 1 h at 37°C. The slices were then incubated in the primary antibodies overnight at 4°C. The primary antibodies, including rabbit anti-ATP8A2 (1:200; Abcam, Cambridge, MA, USA) and mouse anti-NeuN (1:1000; Abcam, Cambridge, MA, USA), were diluted in PBS containing 1% BSA and 0.25% Triton X-100. Afterward, the specimens were washed three times in PBS and then were incubated with secondary antibodies, including Alexa Fluor 555-conjugated goat anti-rabbit and Alexa Fluor 488-conjugated donkey anti-mouse for 2 h at 37°C. Both secondary antibodies (Invitrogen, Thermo Fisher Scientific, Waltham, MA, USA) were diluted to 1:400.

A Stereo Investigator stereological system (MicroBrightField, Williston, USA) was used for quantitative analyses of the ATP8A2^+^ cells in the unilateral PFC of each mouse. Coronal sections of the PFC were collected through the rostrocaudal axis spanning approximately from +3.20 mm to +2.10 mm relative to bregma (Xiong et al.,2017)to count the interested cells. Measurements were recorded from an equidistant series of six coronal sections. After the actual section thickness was measured, appropriate guard zones at the top and bottom of each section were defined to avoid oversampling. The 40× objective of a Nikon microscope was used for all stereological analyses. The numbers of ATP8A2^+^/NeuN^+^ and NeuN^+^ cells within the traced region (PFC) in each of the six selected sections were straightly quantified without using a grid or counting frames. A Zeiss LSM780 confocal laser-scanning microscope (Carl Zeiss AG, Oberkochen, Germany) was used to capture the representative confocal micrographs of the labeled cells.

For ATP8A2 in situ densitometric analysis, one 40-μm-thick coronal section at the middle of the PFC (+2.10 mm relative to bregma) was collected from each animal. Single labeling for ATP8A2 immunofluorescently was performed, rather than double labeling for ATP8A2/NeuN, to minimize the possible interference with ATP8A2 positive immunofluorescence signal intensity during staining practice. A Zeiss LSM780 confocal laser-scanning microscope (Carl Zeiss AG, Oberkochen, Germany) was used to capture the micrographs of the same field as the double-labeled micrographs shown in Fig.6D–F. The ATP8A2 signal was measured as a mean gray value using the software ImageJ and the data were shown in Fig.6 as a relative influence. The procedures were: 1) Image-Type-8bit; 2) Image-Adjust-Threshold; 3) Image-Adjust-Auto Threshold; 4) Analyze-Set Measurements; 5) Analyze-Measure.

### 2.7. Forced swim test (FST)

FST is a standard test used as a screen for measuring depressive-like behaviors in mice. The experiment was carried out on the PND90, the mice were brought to the laboratory one week in advance to let them adapt to the environmental room. In this experiment, mice were placed in a plastic cylinder (40 cm in-depth, 20 cm in diameter and filled with water at 24±°C with the water’s height at 25 cm above from the bottom. Each mouse was put in the plastic cylinder in the same way and forced to swim for 6 min in the plastic cylinder. The first two minutes of mouse behavior was not recorded. The following 4 minutes were recorded by a video tracking system EthoVision (Noldus Information Technology B.V., Wageningen, Netherlands) for the analysis of the total time spent by each animal in staying on the water surface without any struggling or swimming during the test. At the end of the experiment, the hair of the mice was blown dry to prevent them from catching a cold.

### 2.8 Tail-suspension test (TST)

TST is also one of the most commonly used experiments to evaluate depressive-like behavior in rodent models. In the present study, TST was performed on the second day after the FST. During this test, each mouse was held below its tail to the edge of the clip and suspended 60 cm above from the floor. The performances of mice were recorded by a video tracking system EthoVision (Noldus Information Technology B.V., Wageningen, Netherlands) four minutes with a 2-min-gap left prior to the recording. The total time spent by each animal in not struggling during the test was analyzed. After all behavioral tests were done, animals were killed by over-anesthetized with 10% chloral hydrate.

### 2.9 Statistical analyses

The data were statistically analyzed using the SPSS 25.0 statistical software (Chicago, IL, USA). *Pearson*’s correlation analysis was used for data shown in Fig.3. Data shown in Fig.1, Fig.2, Fig.4, Fig.5 and Fig.6 were statistically analyzed using a randomized block design ANOVA to account for the effects of treatment (such as LPS injection and/or anti-IFN-γ neutralizing mAb injection) by modeling both litter factor and sex factor as a fixed effect described as “block effect” in section Results and the treatment factor as a random effect described as “treatment effect” in section Results. For data obtained from more than two treatment groups, the performance of randomized block design ANOVA were followed by Tukey’s *post hoc* test. *p* values are displayed as follows: n.s. = not significant, * *p* < 0.05, ** *p* < 0.01, *** *p* < 0.001.

## 3. Results

### 3.1 Neonatal LPS exposure induced a transiently down-regulated expression of ATP8A2 in the PFC in mice

To determine whether neonatal LPS exposure influence the expression of ATP8A2 in the PFC in mice and how long this potential influence would last following LPS exposure, the first experiment was carried out. Mice in the LPS group were injected with LPS and those in the CON group with PBS of the same volume. Two groups of mice were sacrificed after 1 day, 2 days, 4 days, or 10 days after the last LPS injection. The left unilateral PFC tissue from every mouse was taken on ice to be homogenate immediately for Western blot analyses of ATP8A2 level. The right unilateral PFC tissue was taken on ice and stored at −70°C at once for ELISA analyses of LPS-induced pro-inflammatory cytokines if necessary.

As shown in Fig.1, the ATP8A2 levels significantly decreased at the former three test time points: 1 day (randomly block design ANOVA, block effect: *F_(5,5)_* = 1.120, *p* = 0.452, *n* = 6; treatment effect: *F_(1,5)_* = 44.835, *p* = 0.001, *n* = 6), 2 days (randomly block design ANOVA, block effect: *F_(5,5)_* = 1.369, *p* = 0.369, *n* = 6; treatment effect: *F_(1,5)_* = 102.221, *p* < 0.001, *n* = 6) and 4 days (randomly block design ANOVA, block effect: *F_(5,5)_* = 0.811, *p* = 0.588, *n* = 6; treatment effect: *F_(1,5)_* = 37.424, *p* = 0.002, *n* = 6) after the last LPS injection. However, such decrease was no longer detectable 10 days after the last LPS injection (randomly block design ANOVA, block effect: *F_(5,5)_* = 0.489, *p* = 0.775, *n* = 6; treatment effect: *F_(1,5)_* = 0.078, *p* = 0.791, *n* = 6) (Fig.1). It is worth noting that the largest extent of the decrease in PFC ATP8A2 expression was observed in the sample obtained 48 h after the last LPS injection (Fig.1). These findings verified the expected influence of neonatal LPS exposure on the PFC ATP8A2 expression in mice and revealed the time curve of PFC ATP8A2 expression following neonatal LPS exposure.

**Fig.1.**
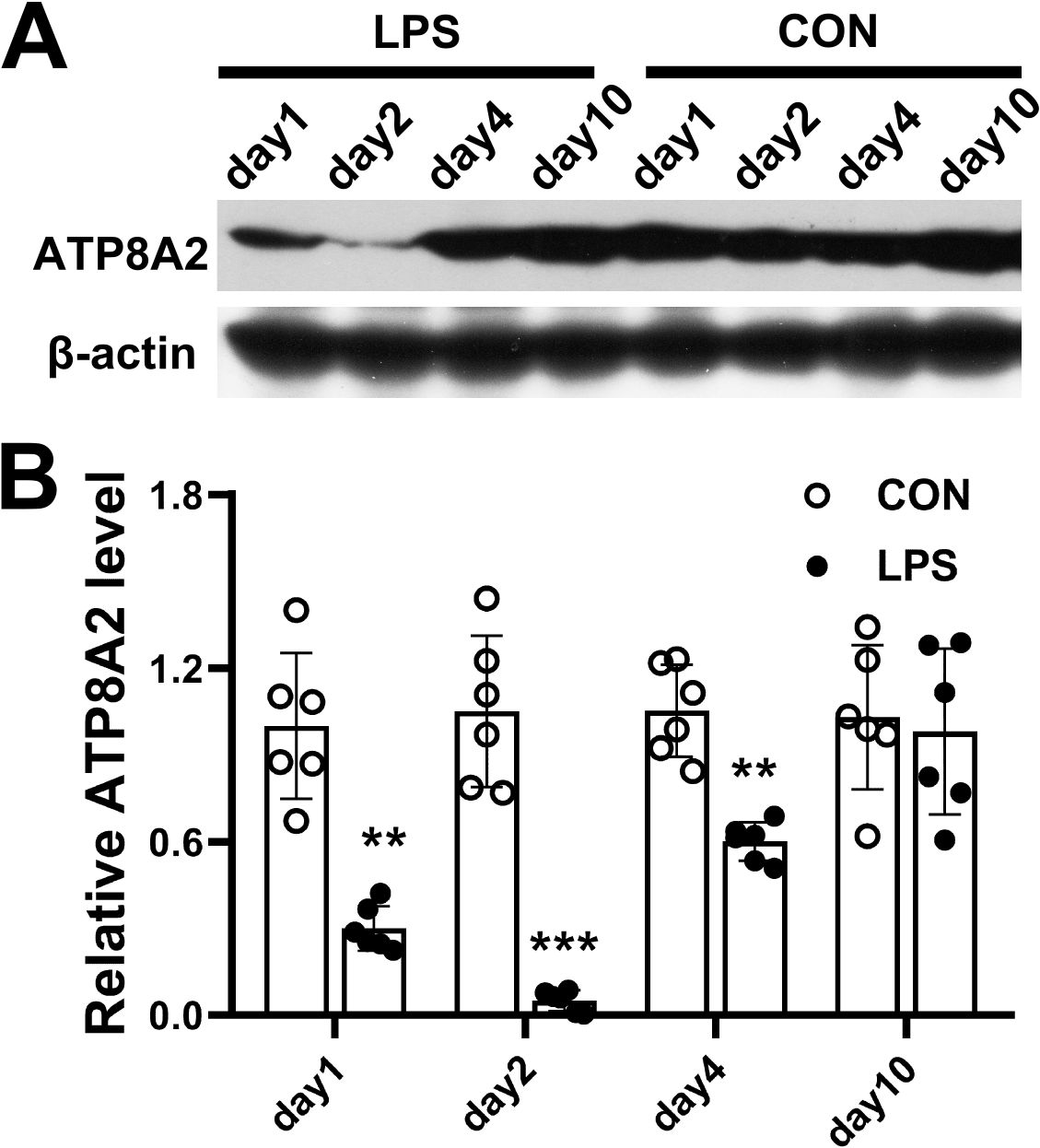
Neonatal LPS exposure induced a transiently down-regulated expression of ATP8A2 in the PFC in mice. **(A)** Representative results for the Western blot analysis of ATP8A2. **(B)** The relative quantification of ATP8A2 in each group of mice was normalized using the level of β-actin. Data are expressed as means ± SEM. Randomly block design ANOVA; *n* = 6/group; * *p* < 0.05; *** *p* < 0.001.

**Fig.2.**
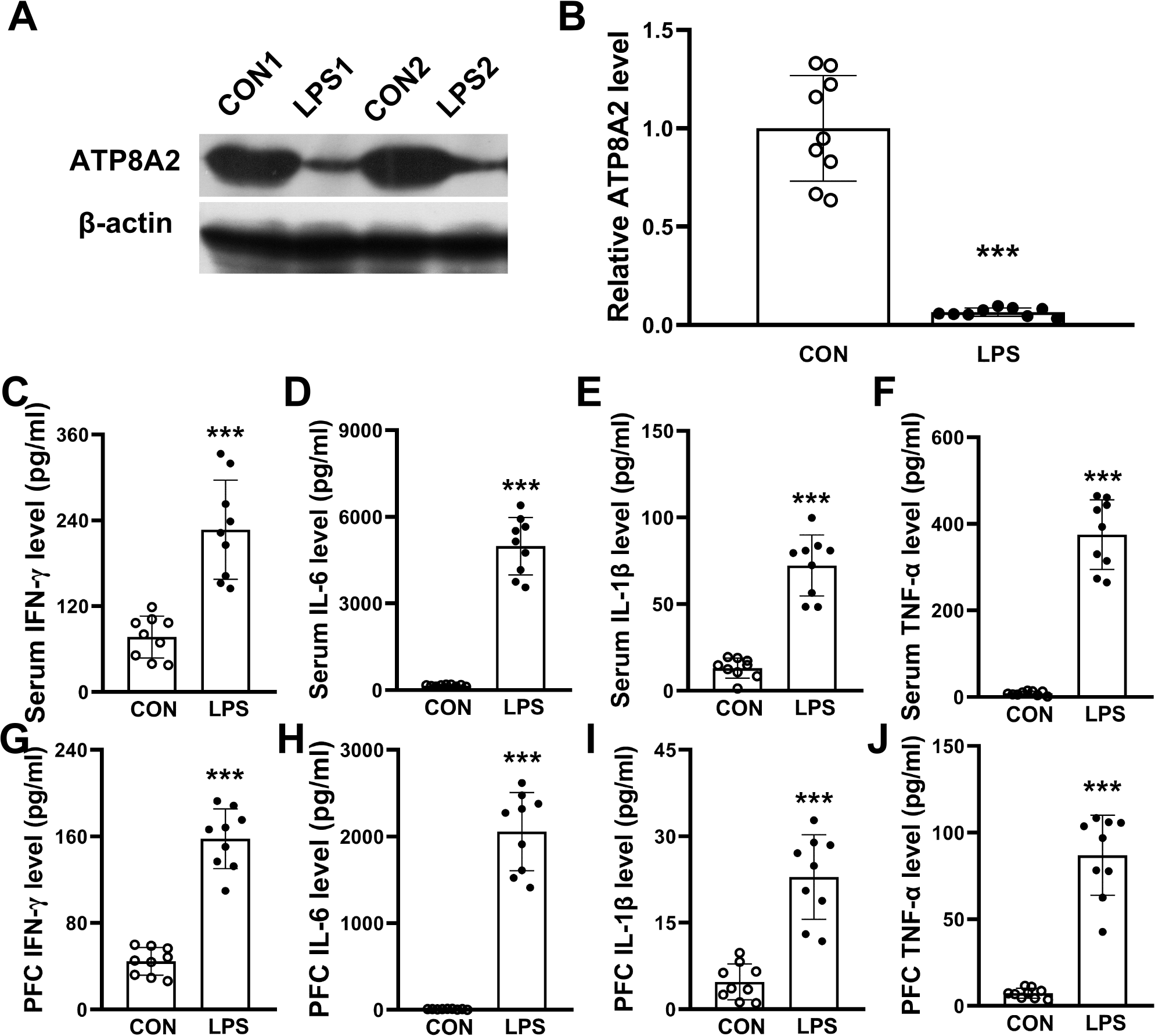
Neonatal LPS exposure induced a down-regulated expression of ATP8A2 accompanied by inflammation both in periphery and in PFC in mice. **(A)** Representative results for the Western blot analysis of ATP8A2. **(B)** The relative quantification of ATP8A2 in each group of mice was normalized using the level ofβ-actin. **(C-J)** The bars represent the average levels of IFN-γ, IL-1β, IL-6, TNF-α in the serum and IFN-γ IL-1β, IL-6, TNF-α in the PFC. Data are expressed as means ± SEM. Randomly block design ANOVA; *n* = 9/group; * *p* < 0.05; \*\**p* < 0.01; *** *p* < 0.001.

### 3.2 PFC ATP8A2 level was only significantly correlated to IFN-γ level among all four elevated pro-inflammatory cytokines both in serum and in the PFC

Given the initial finding that PFC ATP8A2 expression was reduced to the largest extent 48 h after the last LPS injection (Fig.1), the prepared serum and PFC samples (*n* = 6) obtained at the same time point in the last experiment were then subjected to ELISA analyses for the levels of LPS-induced pro-inflammatory cytokines, including IFN-γ, IL-1β, IL-6 and TNF-α.

Moreover, another experiment was performed in which the animal was treated all the same way and the levels of PFC ATP8A2 as well as the levels pro-inflammatory cytokines both in the serum and PFC were detected specifically 48 h after the last LPS injection, using a larger sample size (*n* = 9) so as both to provide a repetitive verification of the very novel finding of ATP8A2 in the brain and to make the total sample size up to *n* = 15 (together with *n* = 6) for this chosen test time point. *Pearson*’s correlation analysis between PFC ATP8A2 level and the level of each of the LPS-induced pro-inflammatory cytokines would get good confidence in the case of *n* = 15.

As shown in Fig.2A–B, the PFC ATP8A2 level decreased by more than 90% in LPS group (randomly block design ANOVA, block effect: *F_(8,8)_* = 1.062, *p* = 0.467, *n* = 9; treatment effect: *F_(1,8)_* = 112.127, *p* < 0.001, *n* = 9). All the detected cytokines, whether in serum (Fig.2C–F) or in PFC (Fig.2G–J), showed a dramatically increased expression (serum IFN-γ: randomly block design ANOVA, block effect: *F_(8,8)_* = 2.310, *p* = 0.129, *n* = 9; treatment effect: *F_(1,8)_* = 68.475, *p* < 0.001, *n* = 9; serum IL-6: randomly block design ANOVA, block effect: *F_(8,8)_* = 0.983, *p* = 0.509, *n* = 9; treatment effect: *F_(1,8)_* = 170.514, *p* < 0.001, *n* = 9; serum IL-1β: randomly block design ANOVA, block effect: *F_(8,8)_* = 1.485, *p* = 0.295, *n* = 9; treatment effect: *F_(1,8)_* = 104.142, *p* < 0.001, *n* = 9. serum TNF-α: randomly block design ANOVA, block effect: *F_(8,8)_* = 0.928, *p* =0.541, *n* = 9; treatment effect: *F_(1,8)_* = 133.012, *p* < 0.001, *n* = 9; PFC IFN-γ: randomly block design ANOVA, block effect: *F_(8,8)_* = 0.722, *p* = 0.672, *n* = 9; treatment effect: *F_(1,8)_* = 107.626, *p* < 0.001, *n* = 9; PFC IL-6: randomly block design ANOVA, block effect: *F_(8,8)_* = 1.001, *p* = 0.499, *n* = 9; treatment effect: *F_(1,8)_* = 129.968, *p* < 0.001, *n* = 9; PFC IL-1β: randomly block design ANOVA, block effect: *F_(8,8)_* = 1.130, *p* = 0.433, *n* = 9; treatment effect: *F_(1,8)_* = 42.304, *p* < 0.001, *n* = 9; PFC TNF-α: randomly block design ANOVA, block effect: *F_(8,8)_* = 1.159, *p* = 0.420, *n* = 9; treatment effect: *F_(1,8)_* = 250.424, *p* < 0.001, *n* = 9)

Taken all the data from 48 h time point samples from LPS-treated mice together, *Pearson*’s correlation analysis was performed between PFC ATP8A2 level and the level of each of the LPS-induced pro-inflammatory cytokines (*n* = 15). There was a significant negative correlation between ATP8A2 expression in PFC and IFN-γ level both in serum (*p* < 0.01, *Pearson*’s correlation analysis, *n* = 15) (Fig.3 A) and in PFC (*p* < 0.001, *Pearson*’s correlation analysis, *n* = 15) (Fig.3E). No significant correlations were found between ATP8A2 level and all the other pro-inflammatory cytokines (all *p* values > 0.05) (Fig.3 B–D, F–H). These results suggest that IFN-γ plays an important role in the PFC ATP8A2 down-regulation caused by neonatal LPS exposure.

**Fig.3.**
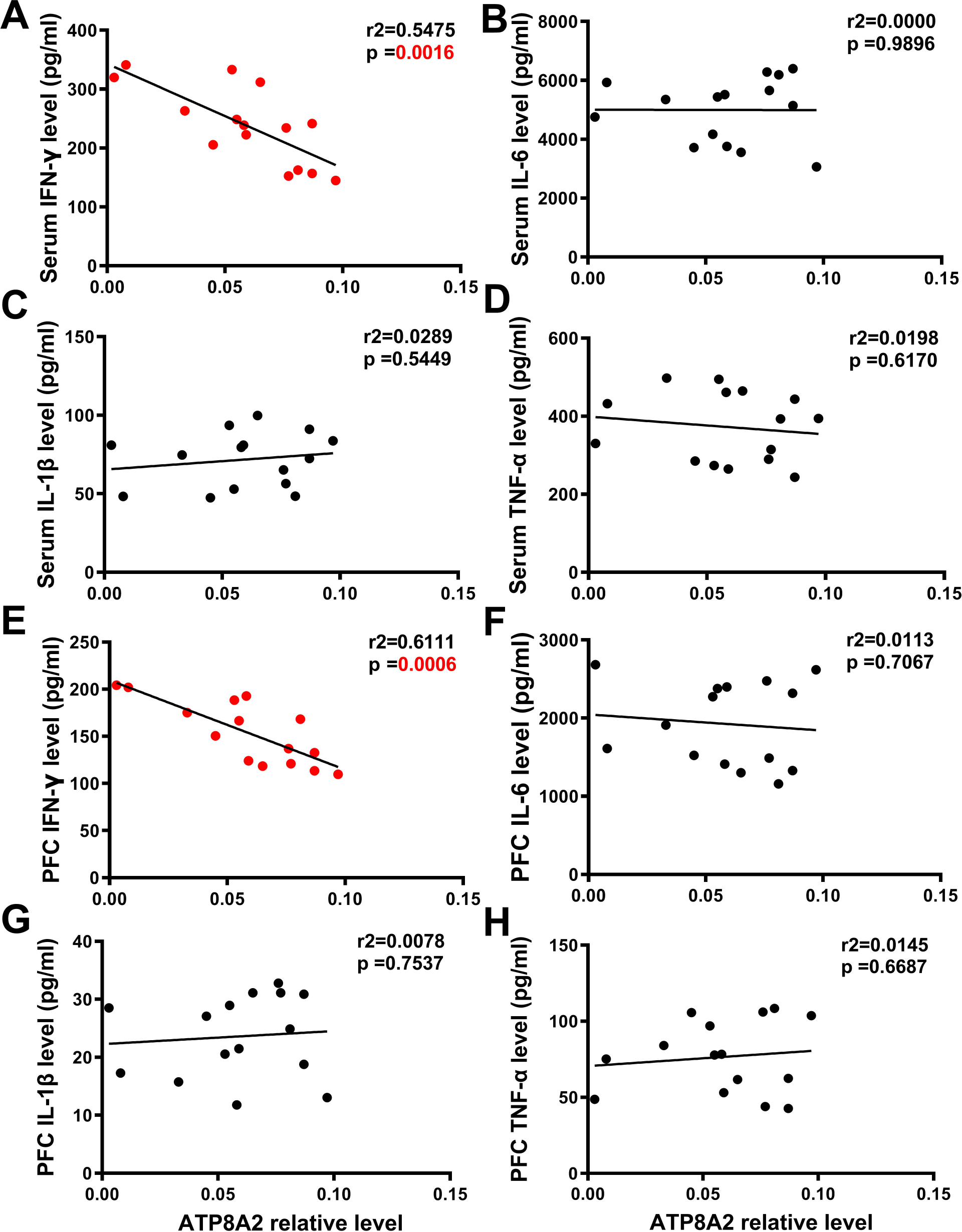
The correlation of PFC ATP8A2 level with the serum and PFC levels of pro-inflammatory cytokines in mice. **(A)** The positive negative correlation of the PFC IFN-γ level with the ATP8A2 expression level in LPS-treated mice. **(B-D)** Correlation analyses between the PFC IL-1β, IL-6β and TNF-α levels and the level of ATP8A2 LPS-treated mice. **(E)** The positive negative correlation of the serum IFN-γ level with the ATP8A2 expression level in LPS-treated mice. **(F-H)** Correlation analyses between the serum IL-1β, IL-6βand TNF-α levels and the level of ATP8A2 LPS-treated mice. *n* = 15 per analysis; *Pearson*’s correlation analysis.

### 3.3 IFN-γ mediated the PFC ATP8A2 down-regulation caused by neonatal LPS exposure

To further explore the potential role of IFN-γ in mediating the PFC ATP8A2 down-regulation following neonatal LPS exposure, an IFN-γ-blocking experiment was conducted using anti-IFN-γ neutralizing mAb and an isotype IgG1. Before this IFN-γ-blocking experiment, another experiment was carried out to determine the optimal dosage of anti-IFN-γ neutralizing mAb. The dosage of 0.6 mg/kg body weight restored the IFN-γγ levels to normal physiological levels both in serum (randomly block design ANOVA, block effect: *F_(7,35)_* = 0.449, *p* = 0.864, *n* = 8; treatment effect: *F_(5,35)_* = 78.511, *p* < 0.001, *n* = 8; *post hoc test*, LPS+anti-IFN-γ (0.6) group vs. CON group: *p* = 1.000) (Fig.4A) and in PFC (randomly block design ANOVA, block effect: *F_(7,35)_* = 0.718, *p* = 0.657, *n* = 8; treatment effect: *F_(5,35)_* = 35.581, *p* < 0.001, *n* = 8; *post hoc test*, LPS+anti-IFN-γ (0.6) group vs. CON group: *p* = 0.821) (Fig.4B). Then, we proceeded to the IFN-γ-blocking experiment and five groups of mice were set. To be specific, the CON group and LPS group were set as above. LPS+anti-IFN-γ (0.6) group was set by injecting LPS combined with anti-IFN-γ neutralizing mAb. The LPS+IgG1 group was set by injecting LPS combined with an isotype IgG1. The last group was set by injecting mere anti-IFN-γ neutralizing mAb (anti-IFN-γ (0.6)).

Randomly block design ANOVA revealed significant differences in IFN-γ levels among five groups (Fig.4C, serum IFN-γ level: randomly block design ANOVA, block effect: *F_(7,28)_* = 0.611, *p* = 0.742, *n* = 8; treatment effect: *F_(4,28)_* = 52.663, *p* < 0.001, *n* = 8) (Fig.4D, PFC IFN-γ level: randomly block design ANOVA, block effect: *F_(7,28)_* = 0.276, *p* = 0.958, *n* = 8; treatment effect: *F_(4,28)_* = 95.873, *p* < 0.001, *n* = 8). We first confirmed that administration of anti-IFN-γ neutralizing mAb blocked the increase of IFN-γ levels both in systemic blood (LPS+anti-IFN-γ (0.6) group vs. CON group: *p* = 0.998) (Fig.4C) and PFC (LPS+anti-IFN-γ (0.6) group vs. CON group: *p* = 0.903) (Fig. 4D), while isotype IgG1 failed to block the LPS-induced IFN-γ increase either in serum (LPS+IgG1 group vs. CON group: *p* < 0.001; LPS+IgG1 group vs. LPS group: *p* = 0.976) (Fig.4C) or in PFC (LPS+IgG1 group vs. CON group: *p* < 0.001; LPS+IgG1 group vs. LPS group: *p* = 0.563) (Fig.4D). In the absence of LPS exposure, the anti-IFN-γ neutralizing mAb significantly neutralized the physiological IFN-γ both in blood (anti-IFN-γ (0.6) group vs. CON group: *p* = 0.013; LPS+IgG1 group vs. LPS group: *p* = 0.976) (Fig.4C) and in brain of mice (anti-IFN-γ (0.6) vs. CON group: *p* < 0.001; LPS+IgG1 group vs. LPS group: *p* = 0.563) (Fig.4D).

**Fig.4.**
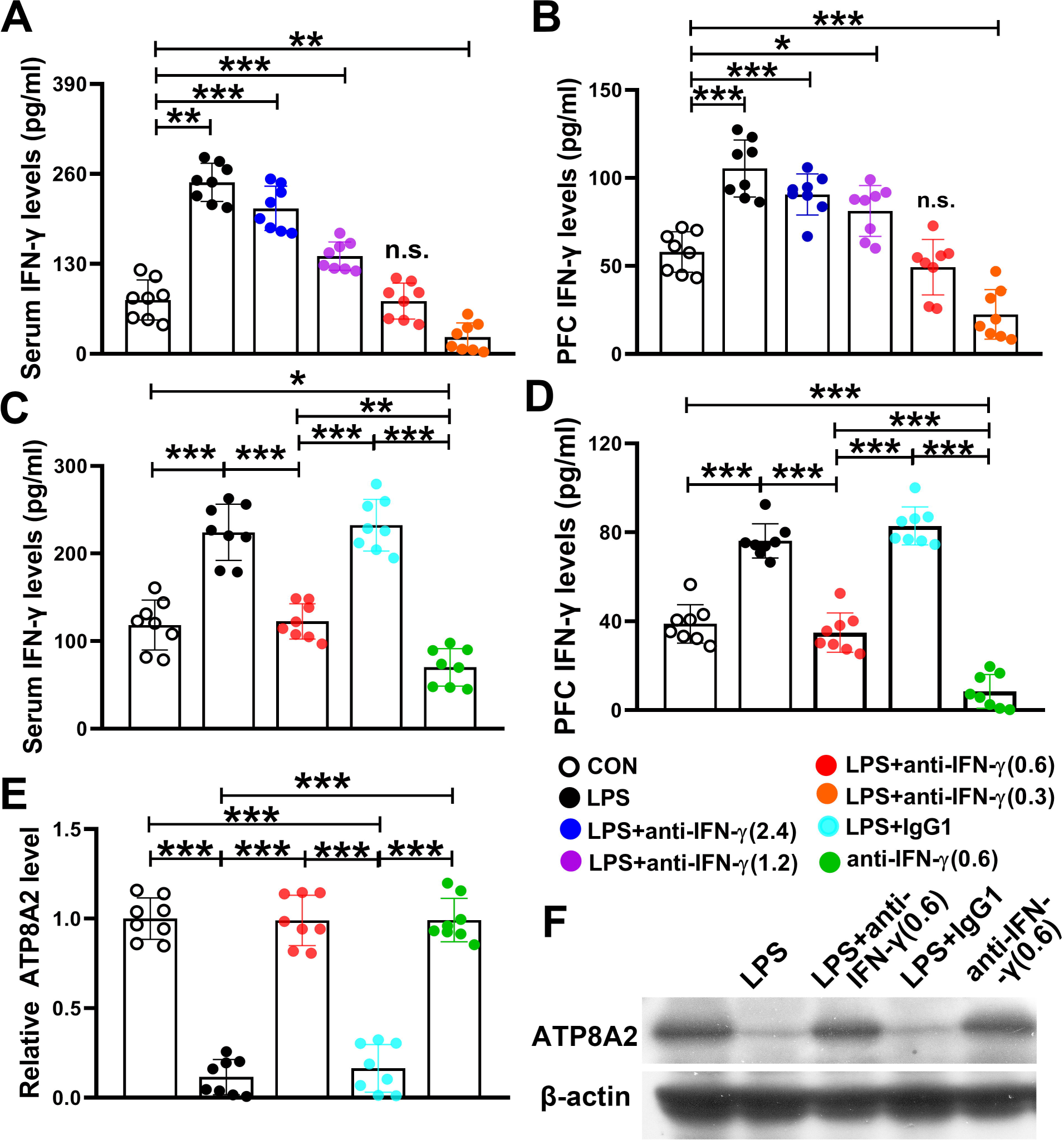
Western blot analyses showed that IFN-γ mediated the PFC ATP8A2 down-regulation caused by neonatal LPS exposure. **(A-B)** The mean levels of IFN-γ in the serum and PFC were shown from the experiment carried out to determine the optimal dosage of anti-IFN-γ neutralizing mAb. **(C-D)** The mean levels of IFN-γ in the serum and PFC were shown from the IFN-γ-blocking experiment. **(E)** The relative quantification of ATP8A2 in each group of mice was normalized using the level of β-actin. **(F)** Representative results for the Western blot analysis of ATP8A2. Data are expressed as means ± SEM. Randomly block design ANOVA followed by Tukey’s *post hoc* test; *n* = 8/group; \*\**p* < 0.01; *** *p* < 0.001; n.s., no significant. anti-IFN-γ (0.6), the anti-IFN-γ dosage of 0.6 mg/kg body weight.

Randomly block design ANOVA of Western blot data revealed significant differences in PFC ATP8A2 levels among five groups (Fig.4E, randomly block design ANOVA, block effect: *F_(7,28)_* = 1.274, *p* = 0.299, *n* = 8; treatment effect: *F_(4,28)_* = 122.680, *p* < 0.001, *n* = 8). We found that neutralization of IFN-γ blocked the LPS-induced ATP8A2 decrease in PFC (LPS+anti-IFN-γ (0.6) group vs. CON group: *p* = 1) (Fig.4E–F), while the isotype IgG1 showed no significant blocking effect on the LPS-induced ATP8A2 decrease in PFC (LPS+IgG1 group vs. LPS group: *p* = 0.922; LPS+IgG1 group vs. LPS+anti-IFN-γ (0.6) group: *p* < 0.001) (Fig.4E–F). Interestingly, in the absence of LPS exposure, mere anti-IFN-γ neutralizing mAb administration led to no significant influence on the PFC ATP8A2 level (anti-IFN-γ (0.6) group vs. CON group: *p* = 1) (Fig.4E–F), although it neutralized the physiological IFN-γ both in the blood (Fig.4C) and in the brain of mice (Fig.4D).

We next performed another experiment to observe the role of IFN-γ using immunofluorescence detection, with a total of three groups set, CON group, LPS group, and LPS+anti-IFN-γ (0.6) group. The groups by giving isotype IgG1 or mere anti-IFN-γ neutralizing mAb were no longer set due to the confirmed results shown above. The results of randomly block design ANOVA of immunofluorescence detection of PFC ATP8A2 levels showed similar findings to Western blot. Neutralization of IFN-γ blocked the LPS-induced ATP8A2 decrease in PFC, indicated by the mean fluorescence intensity (randomly block design ANOVA, block effect: *F_(5,10)_* = 0.473, *p* = 0.788, *n* = 6; treatment effect: *F_(2,10)_* = 47.276, *p* < 0.001, *n* = 6; *post hoc test*, LPS+anti-IFN-γ (0.6) group vs. CON group: *p* = 0.849) (Fig.5K) and ATP8A2^+^/NeuN^+^ cells (randomly block design ANOVA, block effect: *F_(5,10)_* = 0.121, *p* = 0.985, *n* = 6; treatment effect: *F_(2,10)_* = 88.596, *p* < 0.001, *n* = 6; *post hoc test*, LPS+anti-IFN-γ (0.6) group vs. CON group: *p* = 0.532) (Fig.5L). Besides, there were no significant differences among three groups in the number of neurons (NeuN^+^ cells) in PFC (data are not shown) although nearly all patchy ATP8A2^+^ signals were co-located with NeuN^+^ signals (Fig.5G–I). The findings shown in this section identified IFN-γ as the key mediator of the PFC ATP8A2 down-regulation caused by neonatal LPS exposure. Moreover, ATP8A2 alterations were not accompanied by a detectable change in the number of neurons (NeuN^+^ cells) in PFC.

**Fig.5.**
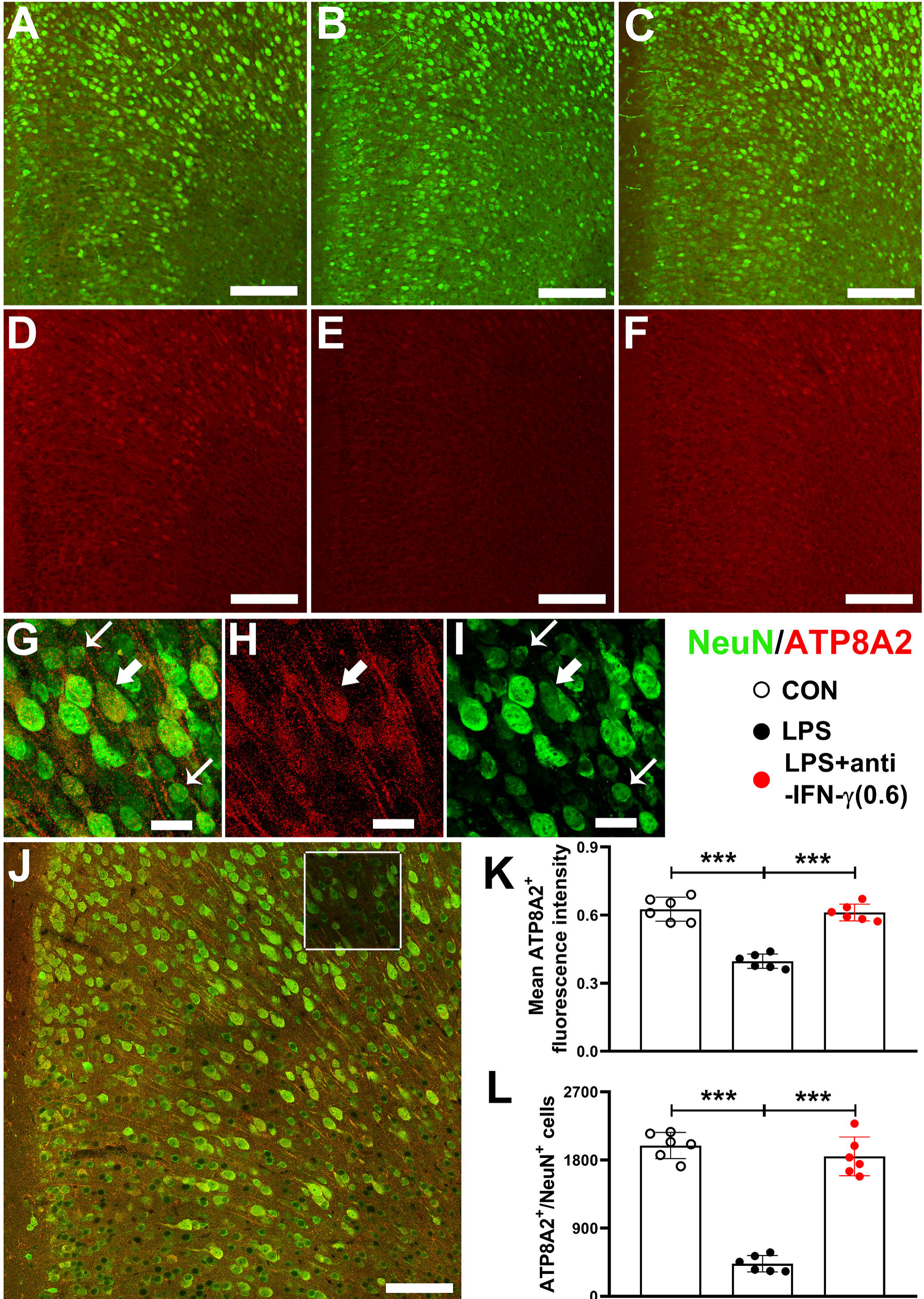
Immunofluorescence analyses showed that IFN-γ mediated the PFC ATP8A2 down-regulation caused by neonatal LPS exposure. **(A-C)** Representative immunofluorescence staining results in the PFC for ATP8A2(red)/NeuN(green) co-labeling in CON group (A), LPS group (B) and LPS+anti-IFN-γ (0.6) group (C). **(E-F)** Exhibition of the single red channel that shows ATP8A2 signal of the same micrographs shown in (A), (B), (C), respectively. **(G-I)** Representative high magnificent photos show the immunofluorescence staining results in the PFC for ATP8A2/NeuN co-labeling. Thick arrows indicate ATP8A2/NeuN co-labeling cells. Thin arrows indicate NeuN single labeling cells. **(J)** The larger scope shows the location in PFC where the photos shown in (G-I) come from. **(K)** Bars represent mean fluorescence intensity of the signal of immunofluorescence staining for ATP8A2 in each group. **(I)** Bars represent mean numbers of ATP8A2/NeuN co-labeling cells in unilateral PFC in each group. Data are expressed as means ± SEM. Randomly block design ANOVA followed by Tukey’s *post hoc* test; *n* = 6/group; *** *p* < 0.001; anti-IFN-γ (0.6), the anti-IFN-γ dosage of 0.6 mg/kg body weight. Scale bar in (A-F), 300 μm; in (G-I), 10 μm; in (J), 200 m.

### 3.4 IFN-γ plays a role in depressive-like behaviors in adulthood induced by neonatal LPS exposure

Abnormality in PFC, especially during the critical development period, is an important pathophysiological basis of depression and it has been verified that *neonatal LPS exposure* could result in depression in adulthood (Dinel et al.,2014; Walker et al.,2013). Therefore, we investigated whether neutralization of IFN-γ affects the behavioral performances in the FST and TST of the LPS-treated mice. The floating immobility time was partially but significantly restored toward the levels of CON group in FST task (randomly block design ANOVA, block effect: *F_(11,44)_* = 0.883, *p* = 0.563, *n* = 12; treatment effect: *F_(4,44)_* = 43.424, *p* < 0.001, *n* = 12; *post hoc test*, LPS+anti-IFN-γ (0.6) group vs. CON group: *p* = 0.002) (Fig.6A). In TST task, neutralization of IFN-γ also resulted in an obvious restoring trend although Tukey *post hoc test* reported the *p* value > 0.05 of the comparision between LPS+anti-IFN-γ (0.6) group and CON group (randomly block design ANOVA, block effect: *F_(11,44)_* = 0.466, *p* = 0.914, *n* = 12; treatment effect: *F_(4,44)_* = 32.685, *p* < 0.001, *n* = 12; *post hoc test*, LPS+anti-IFN-γ (0.6) group vs. CON group: *p* = 0.245) (Fig.6B). However, an LSD *post hoc test* after the same randomly block design ANOVA reported a significant decrease in TST floating immobility time of LPS+anti-IFN-γ (0.6) group than CON group (*p* = 0.043).

The isotype IgG1 failed to block the LPS-induced behavioral alterations either in FST (LPS+IgG1 group vs. CON group: p < 0.001; LPS+IgG1 group vs. LPS group: p = 0.769) (Fig.6A) or in TST (LPS+IgG1 group vs. CON group: *p* < 0.001; LPS+IgG1 group vs. LPS group: *p* = 0.963) (Fig.6B). In the absence of LPS exposure, the anti-IFN-γ neutralizing mAb led to no significant behavioral alterations in FST (anti-IFN-γ (0.6) group vs. CON group: *p* = 0.319; LPS+IgG1 group vs. LPS group: *p* = 0.769) (Fig.6A) or in TST (anti-IFN-γ (0.6) vs. CON group: *p* = 0.999) (Fig.6B). These results suggested a role of IFN-γ in depressive-like behaviors in adulthood induced by neonatal LPS exposure.

### 3.5 No sex dimorphism in ATP8A2 levels or cytokines levels despite a sex dimorphism in depressive-like behaviors in mice

All the analyses described above were intended to exclusively detect the effects of different treatment conditions to test the hypothesis that neonatal LPS exposure might influence PFC ATP8A2 expression in mice involving an increased IFN-γ level and therefore the sex factor was not under consideration as stated in section 2.1. However, previous studies showed that sex may affect immune activation (Cai et al.,2016). So, we also analyzed the data collected from the experiments mentioned in section 3.1 and 3.2 shown in Fig.2 as well as in the bars in Fig.1 indicated as day2 using another statistical method to observe whether there were significant differences in PFC ATP8A2 expression and the levels of proinflammatory cytokines between two sexes either in the presence of LPS challenge or not. The results showed no significant differences in the levels of these molecules between male and female animals either in the presence of LPS challenge or not (all *p* values > 0.05, Supplementary Fig.1). Furthermore, data collected from behavioral tests and shown in section 3.4 (Fig.6) were also subjected to additional analysis for differences between two sexes. The results showed no significant differences in the behavioral task performances of LPS+anti-IFN-γ (0.6) group between male and female animals (FST: LPS+anti-IFN-γ (0.6) group: *p* = 0.536; TST: LPS+anti-IFN-γ (0.6) group: *p* = 0.827; Supplementary Fig.2). However, in all the other groups, significant differences between two sexes were found in FST (CON group: *p* < 0.001; LPS group: *p* < 0.001; LPS+IgG1 group: *p* = 0.018; anti-IFN-γ (0.6) group: *p* = 0.013; Supplementary Fig.2A) and TST (CON group: *p* < 0.001; LPS group: *p* < 0.029; LPS+IgG1 group: *p* < 0.001; anti-IFN-γ (0.6) group: *p* = 0.006; Supplementary Fig.2B) task performances with opposite directions of alterations in the presence of LPS challenge or not. Specifically, female mice showed higher depressive-like behaviors in the absence of LPS challenge, whereas they showed lower depressive-like behaviors in the presence of that.

**Fig.6.**
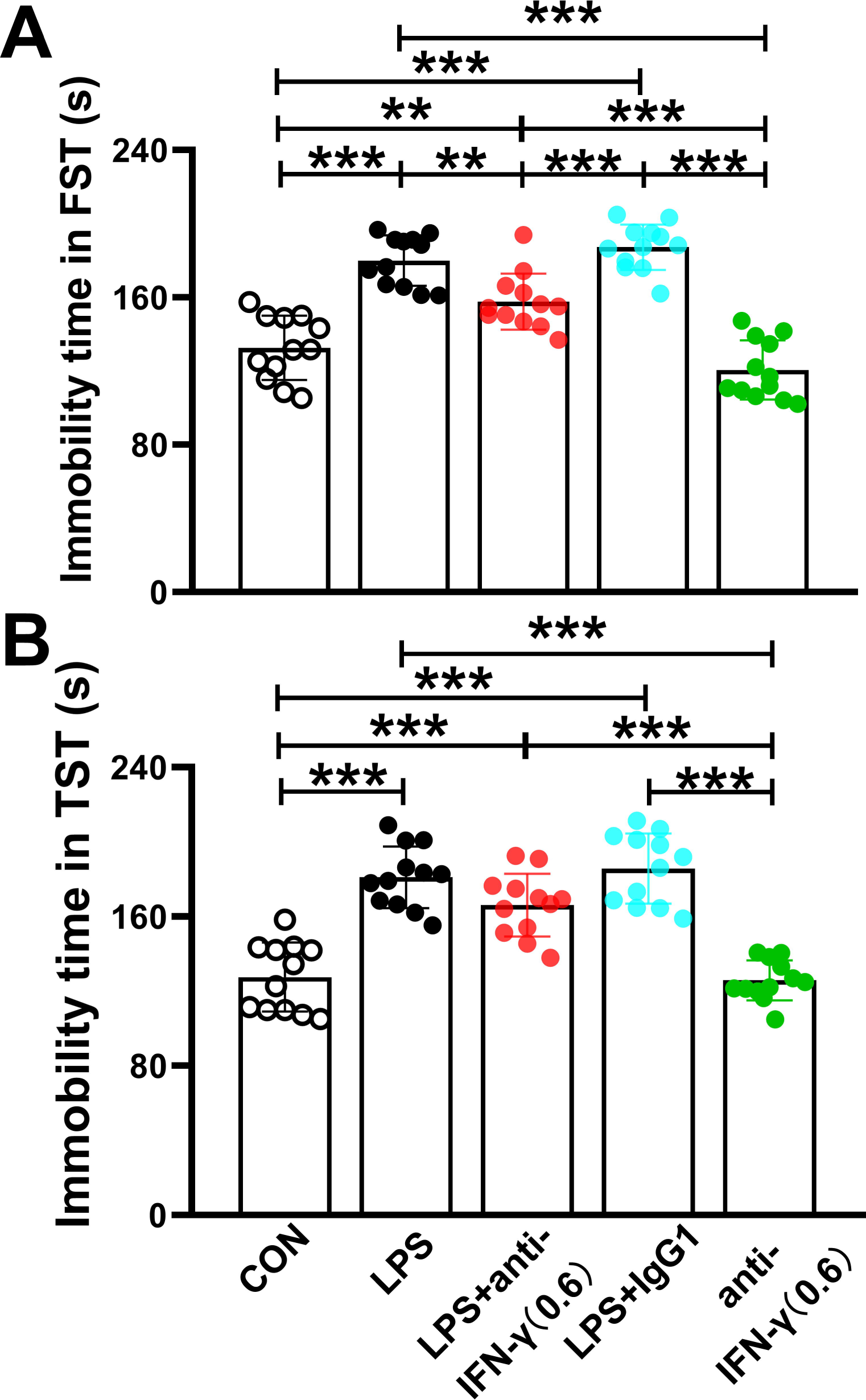
IFN-γ plays an important role in depressive-like behaviors in adulthood induced by neonatal LPS exposure. **(A)** Bars represent the mean immobility time of mice in FST of each group. **(I)** Bars represent the mean immobility time of TST in each group. The data represent the mean ± SEM; randomly block design ANOVA followed by Tukey’s *post hoc* test; *n* = 12/group; \**p* < 0.05, \*\**p* < 0.01, \*\*\**p* < 0.001; anti-IFN-γγ(0.6), the anti-IFN-γ dosage of 0.6 mg/kg body weight.

## Discussion

Our research revealed that neonatal LPS exposure reduced ATP8A2 level in PFC in mice via increasing IFN-γ level. This finding, based on a series of studies concerning the influence of LPS on brain development and behavior (Bilbo and Schwarz,2012; Doosti et al.,2013), is the first to report the change of ATP8A2 in the central nervous system and to further report the mechanism underlying how LPS affects PFC ATP8A2 expression. Moreover, this study further extends our understanding of the mechanism underlying behavioral effects of early life immune activation.

Intraperitoneal LPS injection has been shown to cause a range of acute physiological, pathological, and psychological disorders in rodents. Impairments both in food intake and social exploratory behavior in rodents have been demonstrated in rodents administered intraperitoneally with LPS (Haba et al.,2012; O’Reilly et al.,1988). It has also been reported that LPS exposure can induce depressive and anxiety-like behaviors (Depino,2015; Doosti et al.,2013). Moreover, LPS may reduce acutely the level of prefrontal cortical neurogenesis in adult rodents (Wang et al.,2016). The data from the LPS-treated mice in the present study enriched these widely reported neurobehavioral impairments.

LPS challenge may induce a large extent release of cytokines both in the periphery and brain, particularly pro-inflammatory cytokines, including IFN-γ, IL-1β, IL-6, and TNF-α (Klimstra et al.,1999). These pro-inflammatory cytokines not only play an important role in immunity but also may affect brain function and mediate disease-like behaviors. For example, these pro-inflammatory factors have been shown to cause depressive-like behavior in mice (Gupta et al.,2016; Hashimoto,2015; Kohler et al.,2014; Pokryszko-Dragan et al.,2012). The results of cytokines in our study showed clearly that their levels in the periphery and the brain were highly consistent. this fact may partially due to the immature blood-brain barrier or increased permeability of it during inflammation.

ATP8A2 is a protein located in the membrane, functioning to transport phosphatidylserine into the inner layer of membrane, and thus it is important to maintain the structural stability and normal function of the membrane (Andersen et al.,2016; Coleman et al.,2009). Since the distribution of ATP8A2 in the brain has been determined recently (Andersen et al.,2016), little has been known about the potential factors involved in the regulation of its expression. ATP8A2 expression down-regulation has been observed in certain pathological conditions including Alzheimer’s disease and bacterial infection (Aaron et al.,2018; Ross et al.,2011). However, the study reported a mechanism underlying such regulation could not be seen so far. Decreased level of ATP8A2 in the PFC also occurs in the presence of stress and depressive situation (Chen et al.,2017), but the causal relationship between ATP8A2 change and behavior change is unclear. The current study revealed that ATP8A2 expression in the PFC could be down-regulated by a high concentration of IFN-γ, which is consistent with another study verifying that IFN-γ down-regulated the expression of ATP8A2 in non-neuron cells (Shulzhenko et al.,2018). This finding deepens the understanding of how depression is induced during infections by LPS-producing gram-negative bacteria.

LPS exposure can induce neuroinflammation in various brain regions and result in functional abnormalities, such as hippocampal neuroinflammation and impaired learning and memory (Lee et al.,2008; Shaw et al.,2001). In this study, the animal model that simulates a depressive-like phenotype caused by early bacterial infection was used to investigate whether LPS caused alteration in ATP8A2 expression. Therefore, PFC was selected as the brain region to observe ATP8A2 expression that it’s the area of the brain mostly focused on by studies about depressive-like behavior (Myers-Schulz and Koenigs,2012). In future studies, we will investigate the expression of ATP8A2 in other pathological conditions and/or in other brain zones such as the hippocampus.

As well-known, LPS-induced inflammation is transient. Likewise, the first experiment in this study has shown the ATP8A2 levels decreased transiently and restored within 10 days after the end of LPS administration (Fig.1). Therefore, the animals were killed after finishing behavioral tasks by over-anesthetized without investigating their PFC ATP8A2 levels or peripheral/cerebral cytokines in adulthood. Understandably, a transient ATP8A2 decrease in PFC might mediate a delayed depressive-like behavior phenotype due to the existing theory of early-life programming (Dinel et al.,2014; Karrow,2006), saying that the brain is susceptible to external stimuli, such as immune activation, which modulates the course of normal brain development.

It has been reported that sex could affect the outcomes in behavior development by itself or combining neonatal immune activation (Cai et al.,2016), which is confirmed by the current study due to the observed between-sexes effects in FST and TST tasks performances (Supplementary Fig.2). However, the PFC ATP8A2 expression and cytokine levels both in blood and in PFC have not been significantly influenced by the sex factor tested at PND11 (Supplementary Fig.1). This may be because the sex factor exerts its effects mainly by gonadal hormones. These endocrine sex differences often come to be obvious from the beginning of puberty and these hormonal disparities contribute to the emerging sex differences in the brain (Cai et al.,2016). Further study is required to address whether the sex factor makes a difference in ATP8A2 expression in the brain during puberty as well as adulthood.

In sum, neonatal LPS exposure reduced ATP8A2 level in PFC in mice via increasing IFN-γ level, which may be associated with mechanism underlying brain and behavior impairments induced by neonatal LPS exposure.

## Acknowledgments

We thank Mr. Taoqi Tao (ORCID: 0000-0002-2770-9568, from GDPU), Mrs. Yinyin Xie (ORCID: 0000-0002-5858-3873, from GDPU) for their valuable discussions and help with this investigation.

## Funding Statement

The funders had no role in study design, data collection and interpretation, or the decision to submit the work for publication.

## Funding Information

This paper was supported by the following grants: 1) the National Natural Science Foundation of China (No.31600836) to Junhua Yang; 2) the starting fund for high-level talent introduction into Guangdong Pharmaceutical University (No.51355093) to Junhua Yang; 3) the Innovation and University Promotion Project of Guangdong Pharmaceutical University Through No. 2017KCXTD020 to the School of Biosciences & Biopharmaceutics of GDPU as a team project funding; 4)the National Natural Science Foundation of China (No.81901524) to Li Luo; 5) the Natural Science Foundation of Guangdong Province (No.2018A030313579) to Li Luo.

## Additional information

### Competing interests

JD, LS, ZY, SZ, ZD, LL, JL, XJ and JY declare that there are no conflicts of interest.

### Author contributions

JD: Data curation, Investigation, Methodology, Writing-original draft, Writing-review and editing.

LS: Formal analysis, Investigation, Methodology.

ZY: Data curation, Formal analysis, Investigation, Methodology.

SZ: Validation, Investigation.

ZD: Validation, Investigation.

LL: Resources, Funding acquisition.

JL: Resources.

XJ: Resources.

JY: Conceptualization, Supervision, Funding acquisition, Writing - original draft, Writing - review and editing.

## Notes

### Competing Interest Statement

The authors have declared no competing interest.

