## Supplementary material for "Neonatal LPS exposure reduces ATP8A2 level in the prefrontal cortex in mice via increasing IFN-γ level": Supplementary Material.pdf

**Supplementary figure:**

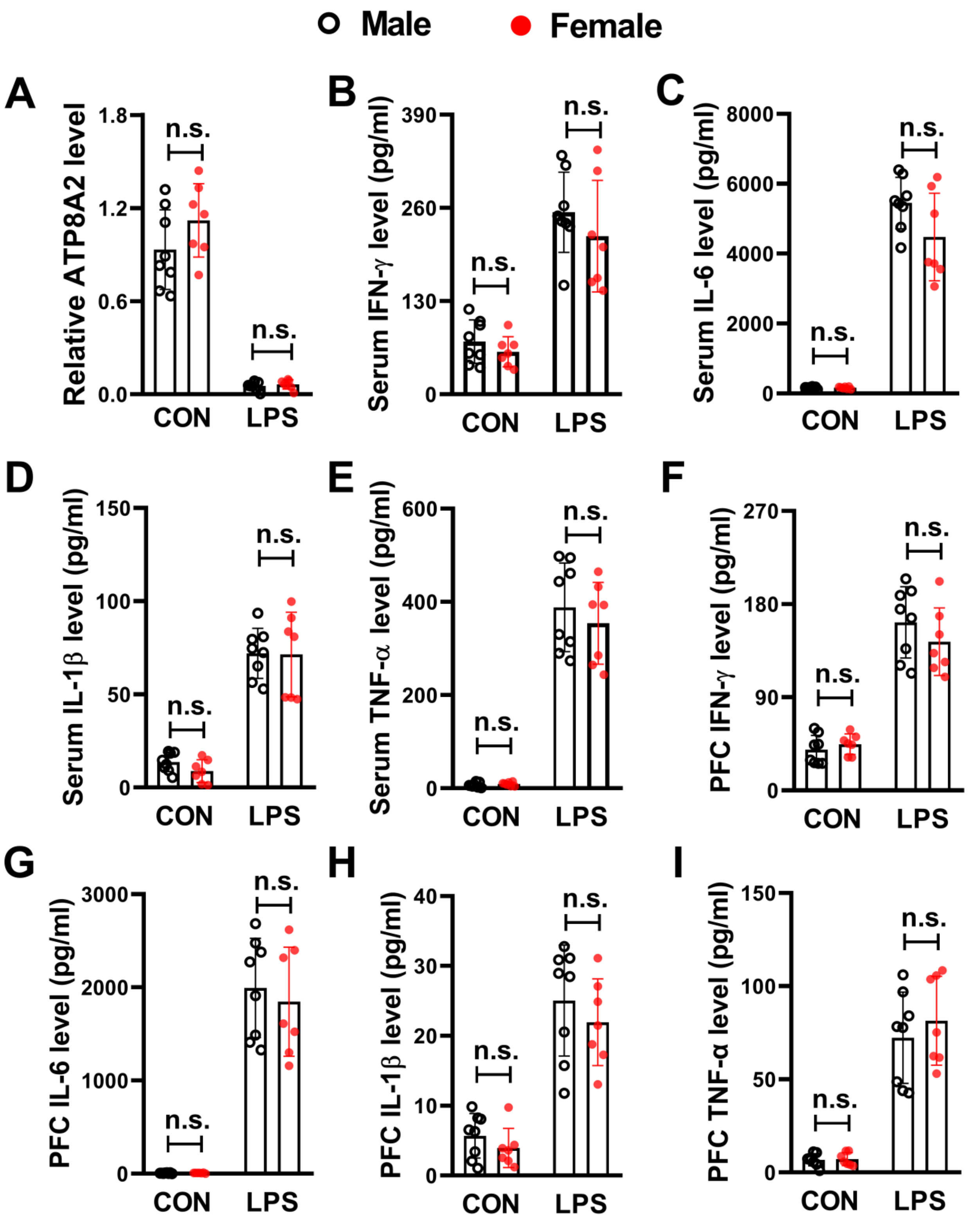


**Supplementary Fig.1. No sex dimorphism in ATP8A2 levels and proinflammatory cytokines levels tested at PND11.** Data shown in this figure were the outputs of statistical analyses of the pooled raw data shown in Fig.2 as well as in the bars in Fig.1 indicated as *day2*, using a different statistical method to explore between-sexes differences. **(A)** Representative results for the Western blot analysis of ATP8A2 proteins in PFC samples. **(B-I)** The bars represent the average levels of IFN-γ, IL-1β, IL-6, TNF-α in the serum (B-E) and those molecules in the PFC (F-I). Welch-*t* test for LPS-treat mice in (C). Mann-Whitney *U* test was performed for data from CON mice shown in (F), (G) and (H) and LPS group shown in (D). The rest data shown in this figure were analyzed using Student's *t* test. The data represent the mean ± SEM. $n$ = 8 per each male group; $n$ = 7 per each female group; n.s., no significant.

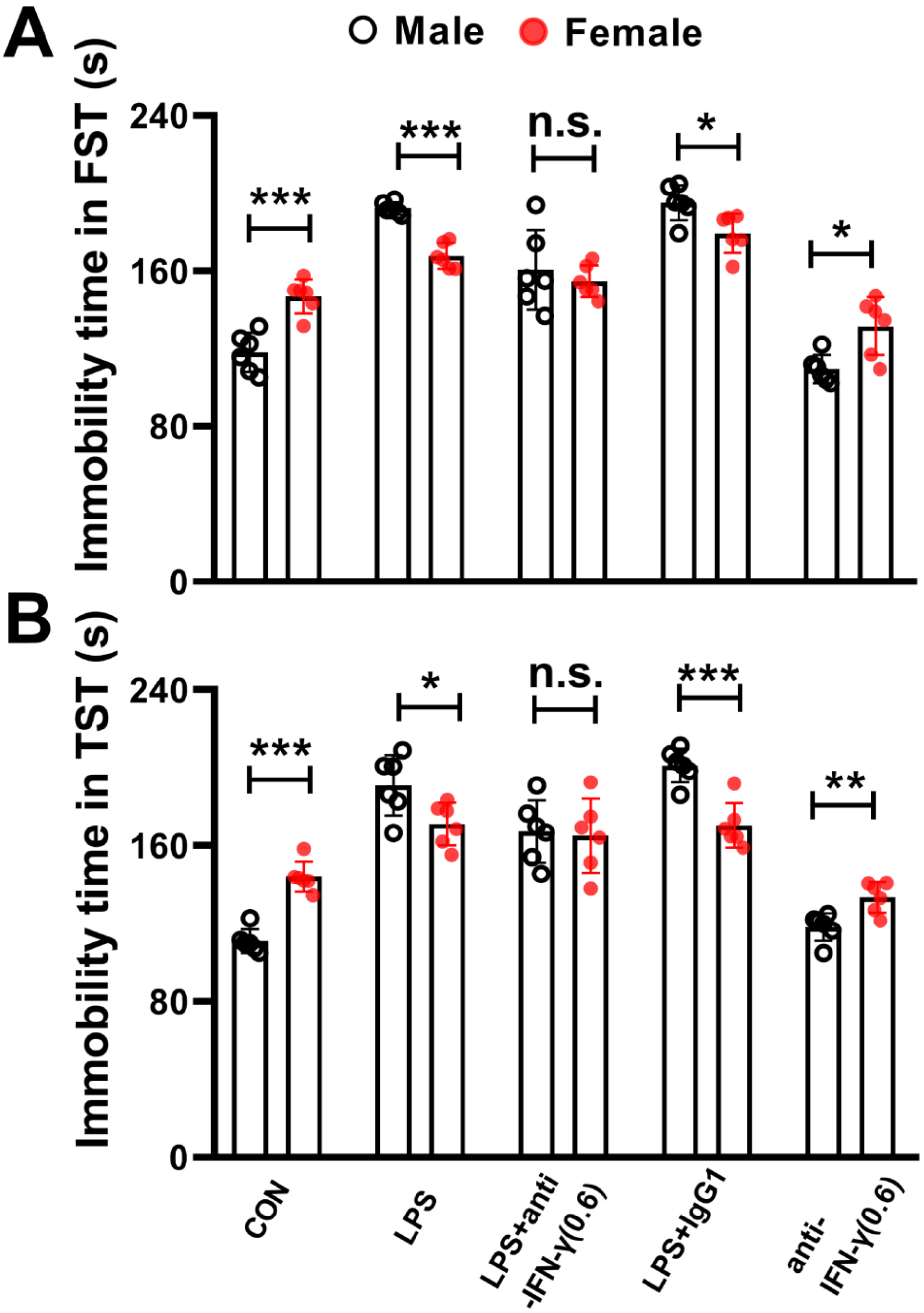


**Supplementary Fig.2. A sex dimorphism in depressive-like behaviors in mice tested at PND90.** Data shown in this figure were the outputs of statistical analyses of the same raw data shown in Fig.6, using a different statistical method to demonstrate between-sexes differences. **(A)** Bars represent the mean immobility time of mice in FST of each group. **(I)** Bars represent the mean immobility time of TST in each group. Welch-*t* test for LPS-treat mice, LPS+anti-IFN-γ(0.6)-treated mice and anti-IFN-γ(0.6)-treated mice in (A). The rest data shown in this figure were analyzed using Student's *t* test. The data represent the mean ± SEM. $n$ = 6 per each male group; $n$ = 6 per each female group; *$p$ < 0.05, **$p$ < 0.01, ***$p$ < 0.001; anti-IFN-γ(0.6), n.s., no significant.

**Supplementary tables:**

**Supplementary table1. Animal information for Fig.1**

| **Litter number** | **Number of pups subjected to analyses in each litter** | **Test time points (**days after the last LPS injections**)** | **Groups** | **Number of pups from each litter divided in each group** | **sex** |
|---|---|---|---|---|---|
| litter1 | 8 | 1 | CON | 1 | m |
| | | | LPS | 1 | m |
| | | 2 | CON | 1 | f |
| | | | LPS | 1 | f |
| | | 4 | CON | 1 | m |
| | | | LPS | 1 | m |
| | | 10 | CON | 1 | f |
| | | | LPS | 1 | f |
| litter2 | 8 | 1 | CON | 1 | f |
| | | | LPS | 1 | f |
| | | 2 | CON | 1 | m |
| | | | LPS | 1 | m |
| | | 4 | CON | 1 | f |
| | | | LPS | 1 | f |
| | | 10 | CON | 1 | m |
| | | | LPS | 1 | m |
| litter3 | 8 | 1 | CON | 1 | m |
| | | | LPS | 1 | m |
| | | 2 | CON | 1 | f |
| | | | LPS | 1 | f |
| | | 4 | CON | 1 | m |
| | | | LPS | 1 | m |
| | | 10 | CON | 1 | f |
| | | | LPS | 1 | f |
| litter4 | 8 | 1 | CON | 1 | f |
| | | | LPS | 1 | f |
| | | 2 | CON | 1 | m |
| | | | LPS | 1 | m |
| | | 4 | CON | 1 | f |
| | | | LPS | 1 | f |
| | | 10 | CON | 1 | m |
| | | | LPS | 1 | m |
| litter5 | 8 | 1 | CON | 1 | m |
| | | | LPS | 1 | m |
| | | 2 | CON | 1 | f |
| | | | LPS | 1 | f |
| | | 4 | CON | 1 | m |
| | | | LPS | 1 | m |

| | | | | | |
|---|---|---|---|---|---|
| | | 10 | CON | 1 | f |
| | | | LPS | 1 | f |
| litter6 | 8 | 1 | CON | 1 | f |
| | | | LPS | 1 | f |
| | | 2 | CON | 1 | m |
| | | | LPS | 1 | m |
| | | 4 | CON | 1 | f |
| | | | LPS | 1 | f |
| | | 10 | CON | 1 | m |
| | | | LPS | 1 | m |

**Supplementary table2. Animal information for Fig.2**

| **Litter number** | **Number of pups subjected to analyses in each litter** | **Test time points (**days after the last LPS injections**)** | **Groups** | **Number of pups from each litter divided in each group** | **sex** |
|---|---|---|---|---|---|
| litter1 | 2 | 2 | CON | 1 | m |
| | | | LPS | 1 | m |
| litter2 | 2 | 2 | CON | 1 | f |
| | | | LPS | 1 | f |
| litter3 | 2 | 2 | CON | 1 | m |
| | | | LPS | 1 | m |
| litter4 | 2 | 2 | CON | 1 | f |
| | | | LPS | 1 | f |
| litter5 | 2 | 2 | CON | 1 | f |
| | | | LPS | 1 | f |
| litter6 | 2 | 2 | CON | 1 | m |
| | | | LPS | 1 | m |
| litter7 | 2 | 2 | CON | 1 | f |
| | | | LPS | 1 | f |
| litter8 | 2 | 2 | CON | 1 | m |
| | | | LPS | 1 | m |
| litter9 | 2 | 2 | CON | 1 | m |
| | | | LPS | 1 | m |

**Supplementary table3. Animal information for Fig.4A-B**

| Litter number | Number of pups subjected to analyses in each litter | Test time points (days after the last LPS injections) | Groups | Number of pups from each litter divided in each group | sex |
|---|---|---|---|---|---|
| litter1 | 6 | 2 | CON | 1 | m |
| | | | LPS | 1 | m |
| | | | LPS+anti-IFN-γ(2.4 mg/kg body weight) | 1 | m |
| | | | LPS+anti-IFN-γ(1.2 mg/kg body weight) | 1 | m |
| | | | LPS+anti-IFN-γ(0.6 mg/kg body weight) | 1 | m |
| | | | LPS+anti-IFN-γ(0.3 mg/kg body weight) | 1 | m |
| litter2 | 6 | 2 | CON | 1 | m |
| | | | LPS | 1 | m |
| | | | LPS+anti-IFN-γ(2.4 mg/kg body weight) | 1 | m |
| | | | LPS+anti-IFN-γ(1.2 mg/kg body weight) | 1 | m |
| | | | LPS+anti-IFN-γ(0.6 mg/kg body weight) | 1 | m |
| | | | LPS+anti-IFN-γ(0.3 mg/kg body weight) | 1 | m |
| litter3 | 6 | 2 | CON | 1 | f |
| | | | LPS | 1 | f |
| | | | LPS+anti-IFN-γ(2.4 mg/kg body weight) | 1 | f |
| | | | LPS+anti-IFN-γ(1.2 mg/kg body weight) | 1 | f |
| | | | LPS+anti-IFN-γ(0.6 mg/kg body | 1 | f |

| | | | | | |
|---|---|---|---|---|---|
| | | | weight) | | |
| | | | LPS+anti-IFN-γ(0.3 mg/kg body weight) | 1 | f |
| litter4 | 6 | 2 | CON | 1 | f |
| | | | LPS | 1 | f |
| | | | LPS+anti-IFN-γ(2.4 mg/kg body weight) | 1 | f |
| | | | LPS+anti-IFN-γ(1.2 mg/kg body weight) | 1 | f |
| | | | LPS+anti-IFN-γ(0.6 mg/kg body weight) | 1 | f |
| | | | LPS+anti-IFN-γ(0.3 mg/kg body weight) | 1 | f |
| litter5 | 6 | 2 | CON | 1 | f |
| | | | LPS | 1 | f |
| | | | LPS+anti-IFN-γ(2.4 mg/kg body weight) | 1 | f |
| | | | LPS+anti-IFN-γ(1.2 mg/kg body weight) | 1 | f |
| | | | LPS+anti-IFN-γ(0.6 mg/kg body weight) | 1 | f |
| | | | LPS+anti-IFN-γ(0.3 mg/kg body weight) | 1 | f |
| litter6 | 6 | 2 | CON | 1 | m |
| | | | LPS | 1 | m |
| | | | LPS+anti-IFN-γ(2.4 mg/kg body weight) | 1 | m |
| | | | LPS+anti-IFN-γ(1.2 mg/kg body weight) | 1 | m |
| | | | LPS+anti-IFN-γ(0.6 mg/kg body weight) | 1 | m |
| | | | LPS+anti-IFN-γ(0.3 mg/kg body weight) | 1 | m |
| litter7 | 6 | 2 | CON | 1 | f |

| | | | | | |
|---|---|---|---|---|---|
| | | | LPS | 1 | f |
| | | | LPS+anti-IFN-γ(2.4 mg/kg body weight) | 1 | f |
| | | | LPS+anti-IFN-γ(1.2 mg/kg body weight) | 1 | f |
| | | | LPS+anti-IFN-γ(0.6 mg/kg body weight) | 1 | f |
| | | | LPS+anti-IFN-γ(0.3 mg/kg body weight) | 1 | f |
| litter8 | 6 | 2 | CON | 1 | m |
| | | | LPS | 1 | m |
| | | | LPS+anti-IFN-γ(2.4 mg/kg body weight) | 1 | m |
| | | | LPS+anti-IFN-γ(1.2 mg/kg body weight) | 1 | m |
| | | | LPS+anti-IFN-γ(0.6 mg/kg body weight) | 1 | m |
| | | | LPS+anti-IFN-γ(0.3 mg/kg body weight) | 1 | m |

**Supplementary table4. Animal information for Fig.4C-F**

| Litter number | Number of pups subjected to analyses in each litter | Test time points (days after the last LPS injections) | Groups | Number of pups from each litter divided in each group | sex |
|---|---|---|---|---|---|
| litter1 | 5 | 2 | CON | 1 | f |
| | | | LPS | 1 | f |
| | | | LPS+anti-IFN-γ(0.6 mg/kg body weight) | 1 | f |
| | | | LPS+IgG1 | 1 | f |
| | | | anti-IFN-γ(0.6 mg/kg body weight) | 1 | f |
| litter2 | 5 | 2 | CON | 1 | f |
| | | | LPS | 1 | f |
| | | | LPS+anti-IFN-γ(0.6 mg/kg body weight) | 1 | f |
| | | | LPS+IgG1 | 1 | f |
| | | | anti-IFN-γ(0.6 mg/kg body weight) | 1 | f |
| litter3 | 5 | 2 | CON | 1 | m |
| | | | LPS | 1 | m |
| | | | LPS+anti-IFN-γ(0.6 mg/kg body weight) | 1 | m |
| | | | LPS+IgG1 | 1 | m |
| | | | anti-IFN-γ(0.6 mg/kg body weight) | 1 | m |
| litter4 | 5 | 2 | CON | 1 | m |
| | | | LPS | 1 | m |
| | | | LPS+anti-IFN-γ(0.6 mg/kg body weight) | 1 | m |
| | | | LPS+IgG1 | 1 | m |
| | | | anti-IFN-γ(0.6 mg/kg body weight) | 1 | m |
| litter5 | 5 | 2 | CON | 1 | m |
| | | | LPS | 1 | m |
| | | | LPS+anti-IFN-γ(0.6 mg/kg body weight) | 1 | m |

| | | | | | |
|---|---|---|---|---|---|
| | | | LPS+IgG1 | 1 | m |
| | | | anti-IFN-γ(0.6 mg/kg body weight) | 1 | m |
| litter6 | 5 | 2 | CON | 1 | f |
| | | | LPS | 1 | f |
| | | | LPS+anti-IFN-γ(0.6 mg/kg body weight) | 1 | f |
| | | | LPS+IgG1 | 1 | f |
| | | | anti-IFN-γ(0.6 mg/kg body weight) | 1 | f |
| litter7 | 5 | 2 | CON | 1 | f |
| | | | LPS | 1 | f |
| | | | LPS+anti-IFN-γ(0.6 mg/kg body weight) | 1 | f |
| | | | LPS+IgG1 | 1 | f |
| | | | anti-IFN-γ(0.6 mg/kg body weight) | 1 | f |
| litter8 | 5 | 2 | CON | 1 | m |
| | | | LPS | 1 | m |
| | | | LPS+anti-IFN-γ(0.6 mg/kg body weight) | 1 | m |
| | | | LPS+IgG1 | 1 | m |
| | | | anti-IFN-γ(0.6 mg/kg body weight) | 1 | m |

**Supplementary table5. Animal information for Fig.5**

| Litter number | Number of pups subjected to analyses in each litter | Test time points (days after the last LPS injections) | Groups | Number of pups from each litter divided in each group | sex |
|---|---|---|---|---|---|
| litter1 | 3 | 2 | CON | 1 | m |
| | | | LPS | 1 | m |
| | | | LPS+anti-IFN-γ(0.6 mg/kg body weight) | 1 | m |
| litter2 | 3 | 2 | CON | 1 | f |
| | | | LPS | 1 | f |
| | | | LPS+anti-IFN-γ(0.6 mg/kg body weight) | 1 | f |
| litter3 | 3 | 2 | CON | 1 | m |
| | | | LPS | 1 | m |
| | | | LPS+anti-IFN-γ(0.6 mg/kg body weight) | 1 | m |
| litter4 | 3 | 2 | CON | 1 | f |
| | | | LPS | 1 | f |
| | | | LPS+anti-IFN-γ(0.6 mg/kg body weight) | 1 | f |
| litter5 | 3 | 2 | CON | 1 | f |
| | | | LPS | 1 | f |
| | | | LPS+anti-IFN-γ(0.6 mg/kg body weight) | 1 | f |
| litter6 | 3 | 2 | CON | 1 | m |
| | | | LPS | 1 | m |
| | | | LPS+anti-IFN-γ(0.6 mg/kg body weight) | 1 | m |

**Supplementary table6. Animal information for Fig.6**

| Litter number | Number of pups subjected to analyses in each litter | Test time points (days after the last LPS injections) | Groups | Number of pups from each litter divided in each group | sex |
|---|---|---|---|---|---|
| litter1 | 5 | 90 | CON | 1 | f |
| | | | LPS | 1 | f |
| | | | LPS+anti-IFN-γ(0.6 mg/kg body weight) | 1 | f |
| | | | LPS+IgG1 | 1 | f |
| | | | anti-IFN-γ(0.6 mg/kg body weight) | 1 | f |
| litter2 | 5 | 90 | CON | 1 | f |
| | | | LPS | 1 | f |
| | | | LPS+anti-IFN-γ(0.6 mg/kg body weight) | 1 | f |
| | | | LPS+IgG1 | 1 | f |
| | | | anti-IFN-γ(0.6 mg/kg body weight) | 1 | f |
| litter3 | 5 | 90 | CON | 1 | m |
| | | | LPS | 1 | m |
| | | | LPS+anti-IFN-γ(0.6 mg/kg body weight) | 1 | m |
| | | | LPS+IgG1 | 1 | m |
| | | | anti-IFN-γ(0.6 mg/kg body weight) | 1 | m |
| litter4 | 5 | 90 | CON | 1 | f |
| | | | LPS | 1 | f |
| | | | LPS+anti-IFN-γ(0.6 mg/kg body weight) | 1 | f |
| | | | LPS+IgG1 | 1 | f |
| | | | anti-IFN-γ(0.6 mg/kg body weight) | 1 | f |
| litter5 | 5 | 90 | CON | 1 | m |
| | | | LPS | 1 | m |
| | | | LPS+anti-IFN-γ(0.6 mg/kg body weight) | 1 | m |
| | | | LPS+IgG1 | 1 | m |
| | | | anti-IFN-γ(0.6 mg/kg body weight) | 1 | m |
| litter6 | 5 | 90 | CON | 1 | f |
| | | | LPS | 1 | f |
| | | | LPS+anti-IFN-γ(0.6 mg/kg body weight) | 1 | f |
| | | | LPS+IgG1 | 1 | f |

| | | | | | |
|---|---|---|---|---|---|
| | | | anti-IFN-γ(0.6 mg/kg body weight) | 1 | f |
| litter7 | 5 | 90 | CON | 1 | f |
| | | | LPS | 1 | f |
| | | | LPS+anti-IFN-γ(0.6 mg/kg body weight) | 1 | f |
| | | | LPS+IgG1 | 1 | f |
| | | | anti-IFN-γ(0.6 mg/kg body weight) | 1 | f |
| litter8 | 5 | 90 | CON | 1 | m |
| | | | LPS | 1 | m |
| | | | LPS+anti-IFN-γ(0.6 mg/kg body weight) | 1 | m |
| | | | LPS+IgG1 | 1 | m |
| | | | anti-IFN-γ(0.6 mg/kg body weight) | 1 | m |
| litter9 | 5 | 90 | CON | 1 | f |
| | | | LPS | 1 | f |
| | | | LPS+anti-IFN-γ(0.6 mg/kg body weight) | 1 | f |
| | | | LPS+IgG1 | 1 | f |
| | | | anti-IFN-γ(0.6 mg/kg body weight) | 1 | f |
| litter10 | 5 | 90 | CON | 1 | m |
| | | | LPS | 1 | m |
| | | | LPS+anti-IFN-γ(0.6 mg/kg body weight) | 1 | m |
| | | | LPS+IgG1 | 1 | m |
| | | | anti-IFN-γ(0.6 mg/kg body weight) | 1 | m |
| litter11 | 5 | 90 | CON | 1 | m |
| | | | LPS | 1 | m |
| | | | LPS+anti-IFN-γ(0.6 mg/kg body weight) | 1 | m |
| | | | LPS+IgG1 | 1 | m |
| | | | anti-IFN-γ(0.6 mg/kg body weight) | 1 | m |
| litter12 | 5 | 90 | CON | 1 | m |
| | | | LPS | 1 | m |
| | | | LPS+anti-IFN-γ(0.6 mg/kg body weight) | 1 | m |
| | | | LPS+IgG1 | 1 | m |
| | | | anti-IFN-γ(0.6 mg/kg body weight) | 1 | m |

**Note:** Each litter consisted of about 6-10 pups. In each litter, the superfluous pups that were not subjected to any treatment in the study, including toe-clipping and injections procedures, were kept simultaneously with pups used so as to avoid possible influences on the pups development of removing the unused pups and the overfeeding in presence of lessened pups number. The superfluous pups in the experiment shown in Supplementary table2 were subjected to simultaneous WB and ELISA analyses for PFC ATP8A2 and cytokines which showed no significant differences compared with CON group (data not shown). The superfluous pups in the rest experiments shown in Supplementary table1, Supplementary table3, Supplementary table4, Supplementary table5 and Supplementary table6 were eventually employed in other use reasonably (for example, used in Physiological Experiment Teaching for undergraduates in our university), approved by the Institutional Animal Ethics Committee of Guangdong Pharmaceutical University.
